# Orientation tuning depends on spatial frequency in mouse visual cortex

**DOI:** 10.1101/069997

**Authors:** Inbal Ayzenshtat, Jesse Jackson, Rafael Yuste

**Affiliations:** NeuroTechnology Center, Department of Biological Sciences, Columbia University, New York, NY 10027

**Keywords:** Visual cortex, Cortical tuning, Orientation selectivity, Spatial Frequency, Two-photon, Calcium imaging, Mouse, Asymmetry, Gabor model

## Abstract

The response properties of neurons to sensory stimuli have been used to identify their receptive fields and functionally map sensory systems. In primary visual cortex, most neurons are selective to a particular orientation and spatial frequency of the visual stimulus. Using two-photon calcium imaging of neuronal populations from the primary visual cortex of mice, we have characterized the response properties of neurons to various orientations and spatial frequencies. Surprisingly, we found that the orientation selectivity of neurons actually depends on the spatial frequency of the stimulus. This dependence can be easily explained if one assumed spatially asymmetric Gabor-type receptive fields. We propose that receptive fields of neurons in layer 2/3 of visual cortex are indeed spatially asymmetric, and that this asymmetry could be used effectively by the visual system to encode natural scenes.

**Significance Statement:** In this manuscript we demonstrate that the orientation selectivity of neurons in primary visual cortex of mouse is highly dependent on the stimulus SF. This dependence is realized quantitatively in a decrease in the selectivity strength of cells in non-optimum SF, and more importantly, it is also evident qualitatively in a shift in the preferred orientation of cells in non-optimum SF. We show that a receptive-field model of a 2D asymmetric Gabor, rather than a symmetric one, can explain this surprising observation. Therefore, we propose that the receptive fields of neurons in layer 2/3 of mouse visual cortex are spatially asymmetric and this asymmetry could be used effectively by the visual system to encode natural scenes.

**Highlights:**

– Orientation selectivity is dependent on spatial frequency.
– Asymmetric Gabor model can explain this dependence.

## Introduction

Neurons in the primary visual cortex (V1) have been traditionally described by their receptive field structure and their response characteristics (Hubel and Wiesel, 1962; Skottun et al., 1991). They are classified into two major groups, simple and complex, and exhibit spatially localized receptive fields that consist of distinct elongated On and Off subfields. In mouse visual cortex ~75% of the orientation selective neurons in layer 2/3 are classified as simple cells, showing response characteristics similar to simple cells in visual cortex of monkeys or cats (Niell and Stryker, 2008).

To capture the selectivity of a neuron to a certain feature of a visual stimulus (e.g., orientation, spatial frequency, size, position, speed, etc.), it is convenient to measure one-dimensional (1D) tuning curves that show the average response of a neuron to a specific feature values. Customary tuning models propose that the response strength of a neuron can be predicted based on the similarity between the optimal stimulus of a neuron and the given stimulus. From the 1D tuning curves, which describe the response behavior to one stimulus feature, one can calculate several parameters such as the preferred stimulus, the strength of the selectivity, or the width of the selectivity that quantifies the neuron’s specificity level. In visual cortex for example, since neurons are highly responsive to lines and edges, such curves, which commonly characterize simple cells in V1, are the orientation- and spatial frequency-tuning curves.

However, reducing the complexity of the RF spatial structure to 1D tuning curves in order to study individual features, comes at a cost of losing information that might be critical for understanding the neuronal representation of sensory information. Here, we measured the population responses of neurons in L2/3 of mice V1 to drifting gratings that varied in both orientation and spatial frequency (SF). We used *in vivo* two-photon Ca^2+^ imaging to measure evoked responses from hundreds of V1 neurons. Accordingly, we calculated a two-dimensional (2D) tuning matrix and studied the relationship between orientation and SF selectivity. Then, we compared orientation tuning-curves in various SFs. First we found that the orientation selectivity of a neuron depends strongly on the stimulus SF such that when we presented gratings with higher or lower SF than the optimum, the orientation selectivity was significantly reduced. In addition to a *quantitative* change in the neurons selectivity strength, we also observed a *qualitative* change in the neuron’s preferred stimulus. As we moved away from the optimal SF, to either lower or higher SF, there was a significant shift in the preferred orientation of the neurons. This dependence between orientation and SF selectivity of cells has been previously observed in the visual cortex of primates and cats (Andrews and Pollen, 1979; Vidyasagar and Siguenza, 1985; Webster and De Valois, 1985; Jones et al., 1987; Zhu et al., 2010).

In order to explain this dependence between orientation selectivity and SF, we used the common Gabor model (Gabor, 1946) to predict the responses of a neuron to various stimuli. A Gabor filter is a Gaussian modulated sinusoid, which well describes the receptive fields of simple cells and successfully models their responses (Marcelja, 1980; Field and Tolhurst, 1986; Jones and Palmer, 1987b). However, the classic Gabor model, even though it succeeds to predict multiple neuronal responses, cannot capture the full variety and complexity of the visual system. And indeed, we found that the classic mathematical description of a 2D symmetric Gabor (with either even- or odd-amplitude symmetry) cannot account for our experimental findings. However, spatially modifying the classic model to introduce a 2D asymmetry by way of tilting the Gabor against its elongated axis generates a fundamental change in the response predictions, which qualitatively explains our experimental observations.

The modified Gabor model presented in this paper can explain the response characteristics of a population of neurons and suggests that the receptive field of many cells in layer 2/3 of visual cortex of mice demonstrates a central asymmetry in its 2D spatial organization.

## Materials and Methods

### Animals

Animal handling and experimentation were carried out in accordance with the US National Institutes of Health and Columbia University institutional animal care guidelines. Animals of both sexes were used and were housed in a temperature-controlled environment on a 12h light-dark cycle. We used a total of 5 mice, either WT or VIP-Cre crossed with LSL-tdTomato (P40-80; The Jackson Laboratory).

### Surgery

The mice were placed on a warming plate (37°C) and anesthetized with isoflurane (initially 2% and reduced to 1-1.5% during surgery) administered via nose cone. A custom-made titanium head-plate was attached to the skull using dental cement. Subsequently, a craniotomy (~1×1 mm) was made over the primary visual cortex (3.5-4.5 mm posterior to Bregma, 2.3-2.7 mm lateral to midline; putative monocular region) using a dental drill (Osada, Inc.). An ophthalmic ointment was applied on both eyes to protect the eyes and prevent dehydration during surgery.

### Dye Loading

For bulk loading of cortical neurons, Oregon Green Bapta-1 AM (OGB-1 AM, Molecular Probes) was first mixed with 4 µl pluronic acid (20% in DMSO) and further diluted in 35 µl dye buffer (150 mM NaCl, 2.5 mM KCl, and 10 mM HEPES (pH 7.4)). 50 µM Sulforhodamine-101 (SR101; Molecular Probes) was added to the solution to label astrocytes (Nimmerjahn et al., 2004). Animals were head-fixed and the dye was slowly pressure-injected into the left visual cortex at a depth of ~130-200 µm below the dura surface (layer 2/3) at an angle of 30° through a patch pipette (outer diameter of ~1-2 µm) using Picospritzer II. 2 to 4 injections were carried out at 10 psi for 8 min each under visual control of two-photon imaging (10× water immersion objective, 0.5 NA Olympus, 850 nm excitation). After dye injections, the exposed cortex was covered with agarose (1.5-2%; Sigma-Aldrich) and a cover glass (World Precision Instruments), to reduce brain motion. Data collection began 60-90 minutes after injections to ensure dye uptake across a large number of cells. During data collection, light anesthesia was maintained by isoflurane (0.8-0.9%) administered via nose cone (KOPF Instruments). Heart rate, respiration and oxygen saturation were monitored throughout the experiments using MouseOx (STARR Life Sciences Corp) and respiration rate was used to monitor and control anesthesia levels.

### Two-Photon Ca^2+^ Imaging

Imaging was performed with a two-photon Moveable Objective Microscope (Sutter Instrument) and a mode-locked dispersion-precompensated Ti:sapphire laser (Chameleon Vision II, Coherent). Frames were scanned through a 20× (0.95 NA, Olympus) or 25× (1.05 NA, Olympus) water immersion objective. Laser intensity was controlled via pockels cell (Conoptics) and ranged between 20-70 mW. Scanning and image acquisition were controlled using Mscan (4.07 frames/sec for 512×512 pixels; Sutter Instrument). OGB-1 fluorescence was excited at 950nm. Fluorescence changes collected with a 20x objective typically varied between 5-50%, and between 10-70% with a 25× objective. Emission was collected using green (535/50 nm) and red (610/75 nm) filters (Chroma) simultaneously on two photomultiplier tubes (PMTs).

### Visual Stimulation

Visual stimuli were generated in MATLAB using Psychophysics toolbox (Brainard, 1997; Pelli, 1997; Kleiner et al., 2007) and displayed on a gamma-corrected LCD monitor (Dell; 19 inches, 60 Hz refresh rate) positioned 15 cm from the contralateral eye, roughly at 45° to the long axis of the animal (spanning ~114° vertical by ~140° horizontal of visual space). The presentation of visual stimuli was synchronized with image acquisition using Mscan (Sutter Instrument) and a routine written in MATLAB, such that each stimulus presentation was triggered on the beginning of frame acquisition. The actual time of stimulus presentation was detected with a silicon photodiode (Hamamatsu) attached to the bottom right corner of the screen.

We presented square-wave drifting gratings (100% contrast) for 670 ms, followed by 3-5 sec of uniform gray background (the mean luminance of the gratings) plus a blank condition. Gratings orientation was perpendicular to the drift direction. Gratings were presented at 12 directions of motion in 30° steps, three combinations of 4 or 5 spatial frequencies: [0.02, 0.03, 0.04, 0.06 cpd] or [0.01 0.02 0.04 0.06 cpd] or [0.01 0.02 0.04 0.08 0.16 cpd] and a temporal frequency of 1.5 Hz. All stimuli were block randomized and repeated 5-10 times and the initial phase of the drifting gratings kept constant across all trials. Stimuli were presented in a pseudorandom order but time courses are shown after sorting (see Fig. 1C).

**Figure 1.**
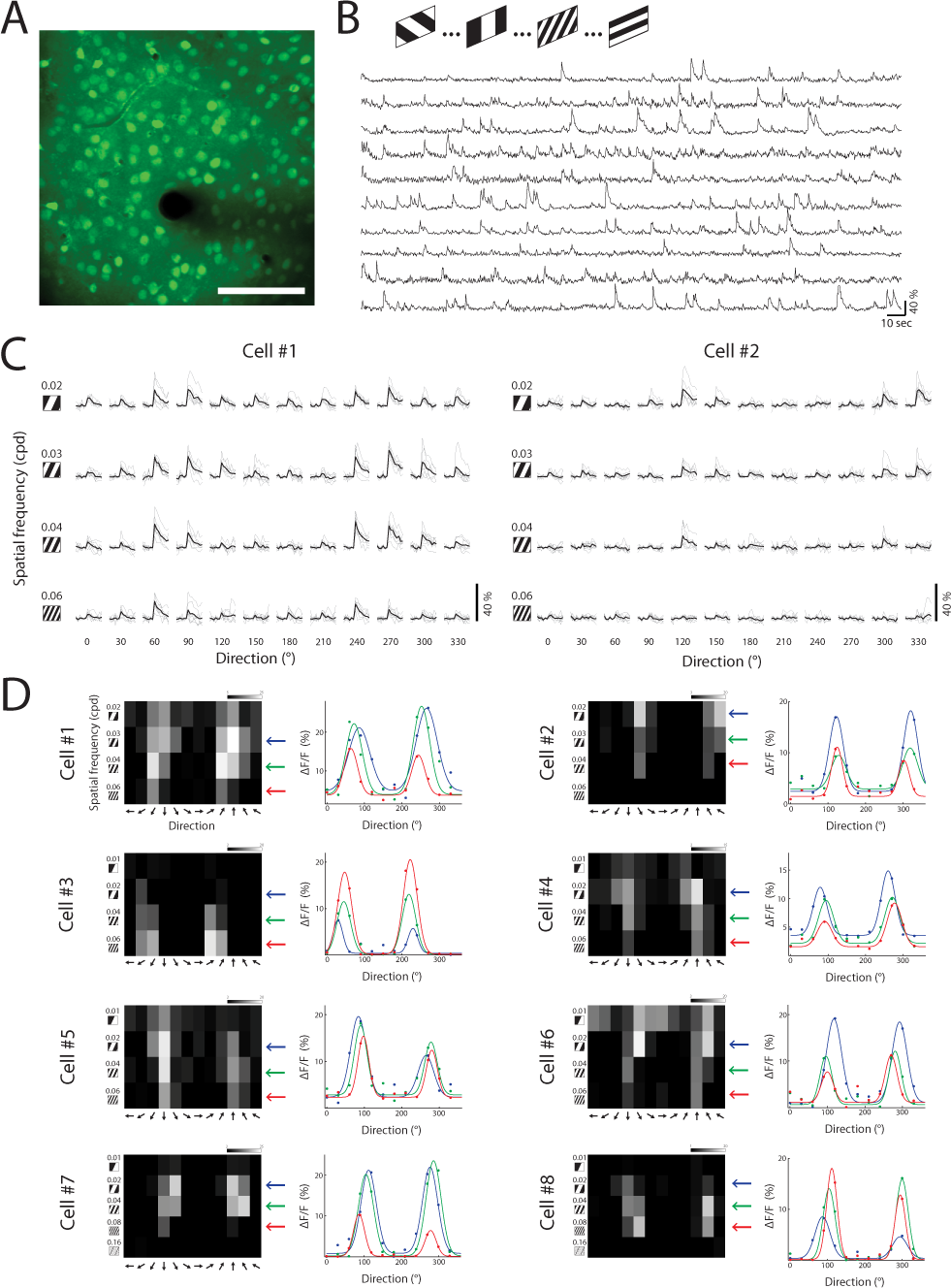
*In vivo* imaging of visual-evoked responses of layer 2/3 neurons. **A,** Top: A two-photon image (maximum projection) from L2/3 neurons in mouse primary visual cortex, loaded with OGB-1. Scale bar, 100 µm. **B,** Example traces of Ca^2+^ signals from 10 cells during the presentation of drifting gratings with 4 different spatial frequencies and 12 directions. **C,** Ca^2+^ responses of two neurons, displayed as a matrix of all stimulus conditions. Columns indicate the direction of motion of the gratings and rows indicate their spatial frequency (SF). Each trial is shown in gray (n=7); average response across trials of a given stimulus is shown in black. **D,** Tuning matrix of 8 cells (cells #1 and #2 are shown in C). Left: response matrices evoked by each stimulus. Pixels intensity corresponds to the average ΔF/F over two frames post stimulus-presentation and over 7 repetitions. Right: Direction-tuning curves fitted with a double Gaussian, measured in various SFs. Colors correspond to SFs marked with arrows on the matrices on the left.

### Data analysis

#### Image Analysis

All data analyses were carried out using built-in and custom built-in software in Matlab (Mathworks). Images were first converted to TIFF format and registered to correct for x-y motion using Turboreg plug-in in ImageJ (Thevenaz et al., 1998). Regions of interest (ROIs) were drawn around each cell using a semi-automated algorithm based on fluorescence intensity (mean projection), florescence change (standard-deviation projections), cell size and shape, and were adjusted by visual inspection. Glia cells were excluded from further analysis using SR101 staining. Pixels were averaged within each ROI for each image frame. Baseline Ca^2+^ fluorescence was computed for each trial as the mean over 2 sec pre-stimulus. Then, fluorescence values were converted to percent change above baseline according to the following: ΔF/F = (F_1_ - F)/F where F is the baseline fluorescence and F_1_ is the instantaneous fluorescence signal averaged over 2 frames (~500 ms) following stimulus onset (F_0+2_,F_0+3_, where F_0_ is stimulus onset frame).

Responsiveness and reliability criteria were defined as previously described (Marshel et al., 2011). Briefly, neurons were considered responsive if their mean ΔF/F to any stimulus exceeds 6%. Reliability (*δ*) was determined according to:

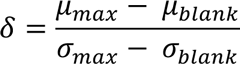

where *µ_max_* and *σ_max_* are the mean and standard deviations of the response to the preferred stimulus respectively, and *µ_blank_* and *σ_blank_* are the mean and standard deviations of the response to the blank stimulus respectively. Neurons were considered reliable if δ > 1. Only cells that demonstrated visual responsiveness and reliability were chosen for further analysis, which excluded between 38 to 43% of the total number of cells we observed per field-of-view (FOV). Therefore we analyzed 85.4±3.6 reliably responsive cells out of 156.2 ± 7.67 cells in total (mean ± SEM; n = 5 mice).

#### Visual Tuning

To calculate tuning curves we averaged the evoked responses (ΔF/F) over 2 frames following stimulus presentation. Then we averaged the response over the number of repetitions (5-10) per stimulus direction (12 directions). Direction tuning curves generated from OGB-1 fluorescence are comparable to those recorded with electrophysiological techniques (Kerlin et al., 2010; Marshel et al., 2011).

Orientation selectivity index (OSI) was computed as follows:

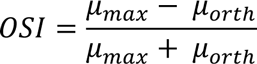

where *µ_max_* is the mean response to the preferred orientation (Pref) and *µ_orth_* is the mean response to the orthogonal orientation (Orth; average of both directions). Only cells that demonstrated OSI ≥ 0.3 were chosen for tuning comparisons.

Orientation tuning curves were fitted with the sum of two Gaussians of identical width, as follows:

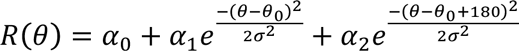

where *R(θ)* is the averaged response to gratings with orientation *θ, α*_0_ is the mean ΔF/F of the 4 lowest points in the curve. *α*_1_ and *α*_2_ are the amplitudes of the two Gaussians, *θ*_0_ is the preferred orientation and *σ* is the standard deviation of the Gaussian function. The sum of two Gaussians fitting was constrained according to Mazurek et al., 2014 using several initial conditions, considering the fit with the lowest least square error as the best fit of the data. Peak responses were the maximum ΔF/F values of the 2D Gaussian fit curve. Half-width at half-height was computed as follows:

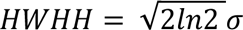

where *σ* is the standard deviation of the Gaussian function.

To further assess the statistical robustness of the tuning curve fitting, we applied a bootstrap method where we randomly resampled the data with replacement for each cell and obtained a distribution of response values. This procedure was repeated 100 times where each repetition was fitted with a double Gaussian (Mazurek et al., 2014). This yielded a distribution of values for each of the tuning curve parameters. The mean values of the resampled data were then used for comparing population’s parameters showing the same statistical significance as comparing parameters of tuning curves obtained by fitting the data that included all the trials.

Additionally, orientation selectivity was assessed using a metric based on 1 minus circular variance (Ringach et al., 2002) as follows:

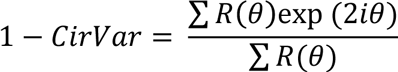

where *R(θ)* is the averaged response to gratings with orientation *θ*. This metric takes into account both tuning width and depth of modulation and found to be more reliable than extracting parameters following curve fitting and more sensitive for detecting differences in selectivity between two populations (Piscopo et al., 2013; Mazurek et al., 2014).

#### 2D Gabor Model

The two-dimensional Gabor function is the product of a 2D sinusoid wave with a circular Gaussian envelope (see Fig. 3A), defined as follows:

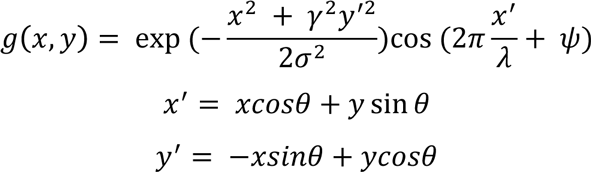

where *σ* is the standard deviation of the Gaussian envelope, which determines the size of the receptive field; *γ* is the spatial aspect ratio of the Gaussian, which determines the ellipticity of the Gabor (in a circular Gaussian *γ* = 1); *λ* is the wavelength of the sinusoid; *θ* is the orientation; and *Ψ* is the phase offset.

**Figure 3:**
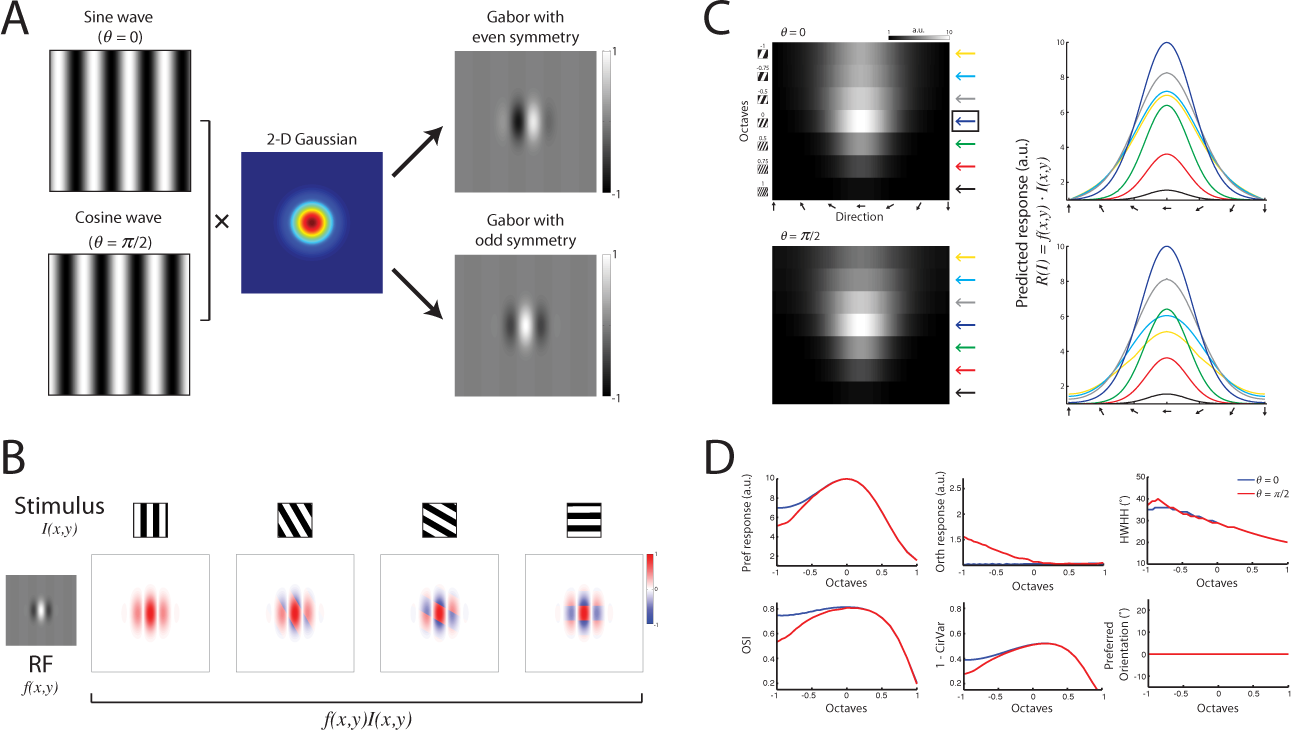
Predicting the responses of a simple cell based on RF-model of 2D symmetric Gabor. **A,** A 2D oriented Gabor function: a sinusoidal plane wave weighted by a Gaussian envelope in two different phases: *θ* = 0 (sine wave) or *θ* = π/2 (cosine wave) shown at the top and bottom panels, that generate Gabor filters with even and odd symmetry, respectively. **B,** Constructing spatiotemporally oriented impulse responses from gratings stimuli drifting against a Gabor RF. Top: examples of stimuli with four different orientations. Bottom: left shows an example of a Gabor RF with *θ* = π/2. On the right, 4 examples where each box depicts the overlap of the grating stimulus crossing the RF in a phase that yields the maximum response, calculated as the inner product between the stimulus and the RF. **C,** A predicted-response tuning matrix computed as the inner product between the Gabor RF model shown in A, and stimuli of square-wave gratings with various orientations (1° interval) and 7 SFs. Each pixel represents the inner product between the RF and the stimulus (maximized across phase, see Materials and Methods). The row marked with a blue arrow on the right denotes the preferred SF and was taken as a reference for comparing other SFs. Right: orientation tuning curves at various SFs color-coded according to the arrows shown next to the predicted tuning matrix. Top and bottom panels correspond to RF with *θ* = 0 and *θ* = π/2, respectively. **D,** Comparison of orientation tuning parameters between various SFs, based on the predicted response shown in C. Blue and red lines depict the parameters calculated based on a Gabor RF model with *θ* = 0 and *θ* = π/2, respectively. Shown are the response amplitude (ΔF/F) to the preferred orientation (Pref), the response amplitude to the orthogonal orientation (Orth), OSI, 1-CirVar, HWHH and the preferred orientation.

In the tilted Gabor model, we introduced another parameter, which is the angle of the Gaussian tilt *(ϕ)* with respect to the sinusoid wave. To generate a tilted Gabor filter, we first generated a symmetric 2D Gaussian with a spatial aspect ratio of *γ* = 0.5 (see also Fig. 4B) and then applied a tilt by multiplying it with a 2D rotation matrix *A* with *ϕ* = 30°:

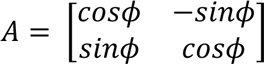

**Figure 4:**
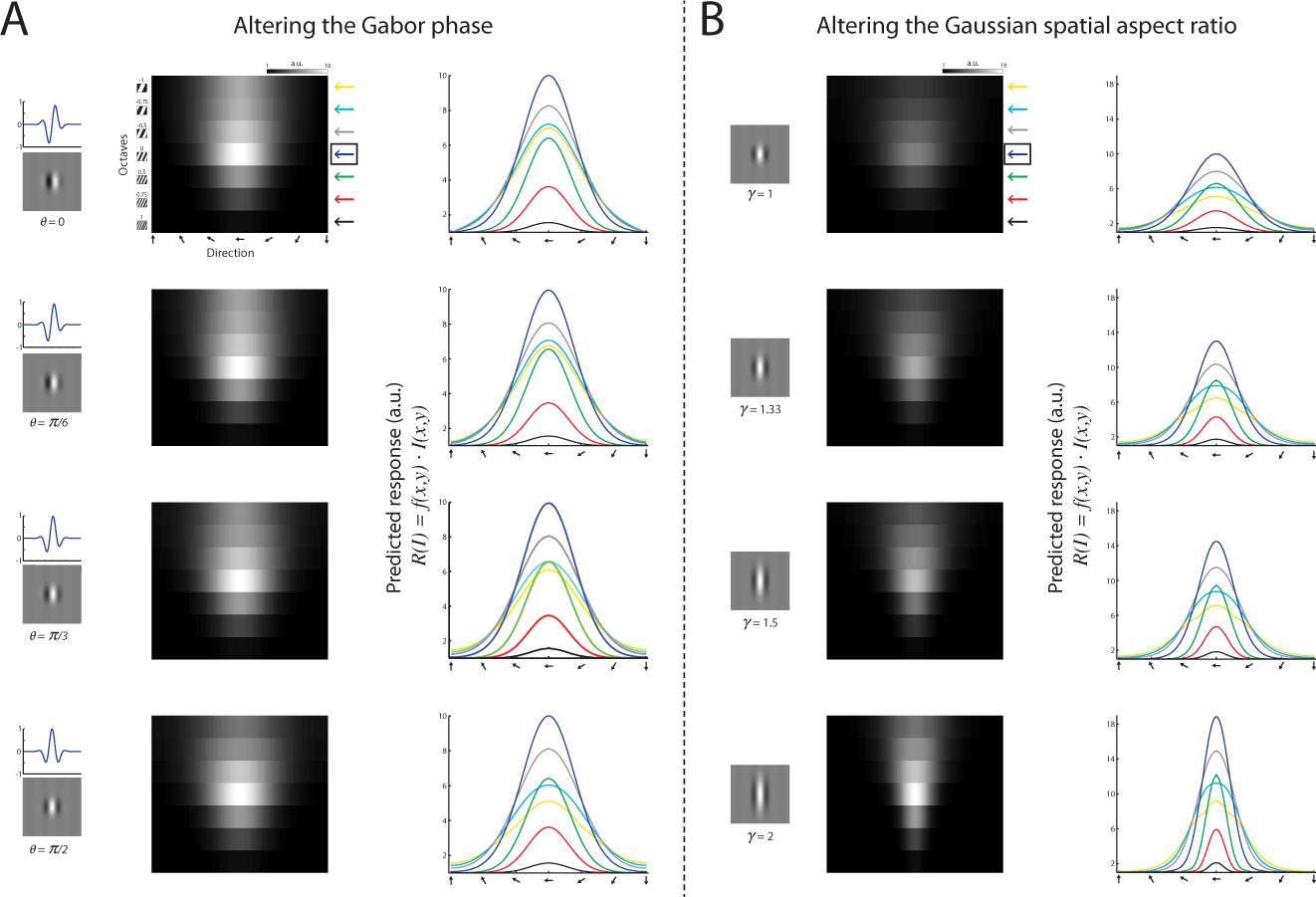
Altering the Gabor model parameters for predicting neural responses. **A**, A predicted-response tuning matrix computed as the inner product between the Gabor RF model shown in Fig. 3 and stimuli of square-wave gratings with various orientations (1° interval) and 7 SFs. Each pixel represents the inner product between the RF and the stimulus (maximized across phase). The row marked with a blue arrow on the right denotes the preferred SF and was taken as a reference for comparing other SFs. Right: orientation tuning curves at various SFs, color-coded according to the arrows shown next to the predicted tuning matrix. Each panel corresponds to RF with *θ* = 0, *θ* = π/6, *θ* = π/3, and *θ* = π/2. **B,** Same as in A only for an RF with *θ* = π/2 but different spatial aspect ratio *(γ)*. Each panel corresponds to RF with *γ* = 1, *γ* = 1.33, *γ* = 1.5, and *γ* = 2. Note that these parameter alterations alone could not qualitatively explain a shift in the preferred orientation of the cells at different SFs.

#### Calculating the predicted response

To calculate the predicted response *R(I)* of a neuron, we computed the inner product between the RF *f(x,y)* - either a symmetric or a tilted Gabor, and the stimulus *I(x,y)* - square-wave gratings drifting in 180 different orientations with 1° interval (see examples in Fig. 3B). Since we measured neuronal responses by recording Ca^2+^ signals, we obtained tuning matrices by maximizing the averaged evoked response per stimulus. Therefore, to compute the predicted response, we calculated the inner product per stimulus orientation for each phase of the stimulus between 0 and 2*π* (21 samples, *π*/10 apart; see Fig. 5) and then maximized the response across phase.

**Figure 5:**
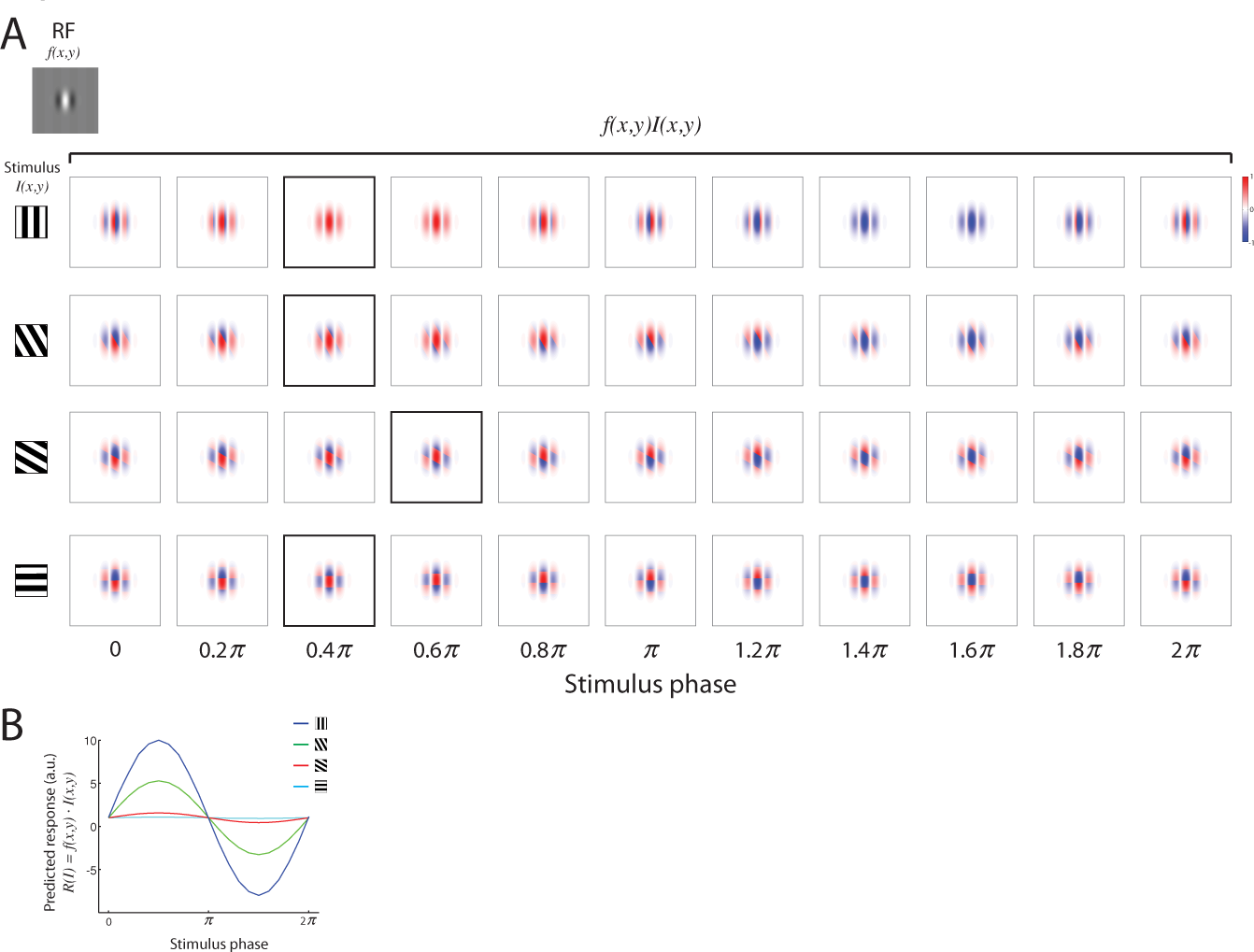
Impulse responses from gratings stimuli drifting against a Gabor RF. **A,** The impulse responses from gratings stimuli drifting against a Gabor RF in various offsets. Top left: a Gabor RF with *θ* = π/2. In each row, on the left is the stimulus presented in one orientation; on the right are 11 examples where each box depicts the overlap of the grating stimulus crossing the RF in a particular phase. The box marked with a black border is the phase yielding the maximum response. **B,** Bottom: The predicted response as a function of phase, calculated as the inner product between the RF and the stimulus at each phase. The predicted response per stimulus was then maximized across phase.

### Results

**Layer 2/3 cells in mouse V1 show dependence of orientation selectivity on spatial frequency**

In lightly anesthetized mice, we identified cells in layer 2/3 of visual cortex using two-photon Ca^2+^ imaging and monitored the activity of neuronal populations (Marshel et al., 2011; Miller et al., 2014) evoked by a brief visual stimulus presented to the contralateral visual field. We characterized the response dynamics by optically recording Ca^2+^ signals of OGB-1 from cells in upper layer 2/3 (~130-200 µm depth, n = 5 animals) in a typical field of view (FOV; Figure 1A-B). Specifically, we measured spatial-frequency- and direction-tuning curves of each neuron in our FOV by averaging across trials the response to each stimulus (Fig. 1C-D). Stimuli were square-wave drifting gratings presented at 1.5 Hz for 670 ms (~ I cycle) and varied across 12 directions and 4 or 5 spatial frequencies (SFs; see Materials and Methods).

Initial characterization of the stimulus-evoked responses of the local network showed dependence of the cells’ orientation selectivity on SF. We characterized this dependence by calculating orientation tuning curves for each SF and then estimating six parameters derived from these curves: a) response amplitude of the preferred orientation (Pref); b) response amplitude of the orthogonal orientation (Orth); c) orientation selectivity index (OSI); d) 1 minus circular variance (1-CirVar); e) half-width at half-height (HWHH); and f) the preferred orientation (see Materials and Methods). For each cell (85.4±3.6 cells/animal, mean±SEM) we compared its tuning curve at the preferred SF, with the tuning curves at SFs below (Fig. 2A) and above (Fig. 2B) the preferred SF.

Tuning curves at SF *below* the preferred SF had by definition reduced responses to the preferred orientation (*p* < 10^−10^, n = 67 cells; Wilcoxon signed-rank test), but also showed increased responses to the orthogonal-to-preferred orientation (*p* < 0.001, Wilcoxon signed-rank test), a decrease in OSI (*p* < 10^−7^, Wilcoxon signed-rank test), a decrease in 1-CirVar (*p* < 10^−6^, Wilcoxon signed-rank test) and an increase in HWHH (*p* < 10^−4^, Wilcoxon signed-rank test) (Fig. 2A). In addition, we observed that some cells had a significant shift in their preferred orientation (Fig 2A, right panel; since we sampled 12 directions between 0° to 360°, for this analysis we subtracted 180° from the preferred direction of cells that had preferred direction > 180°). To quantify this shift regardless of its sign, we calculated the absolute value of the Δ in preferred orientation between the preferred-SF and SF below the preferred. We observed a mean shift of a 22.9 ± 2.8° (mean ± SEM, Fig. 2C left panel).

**Figure 2:**
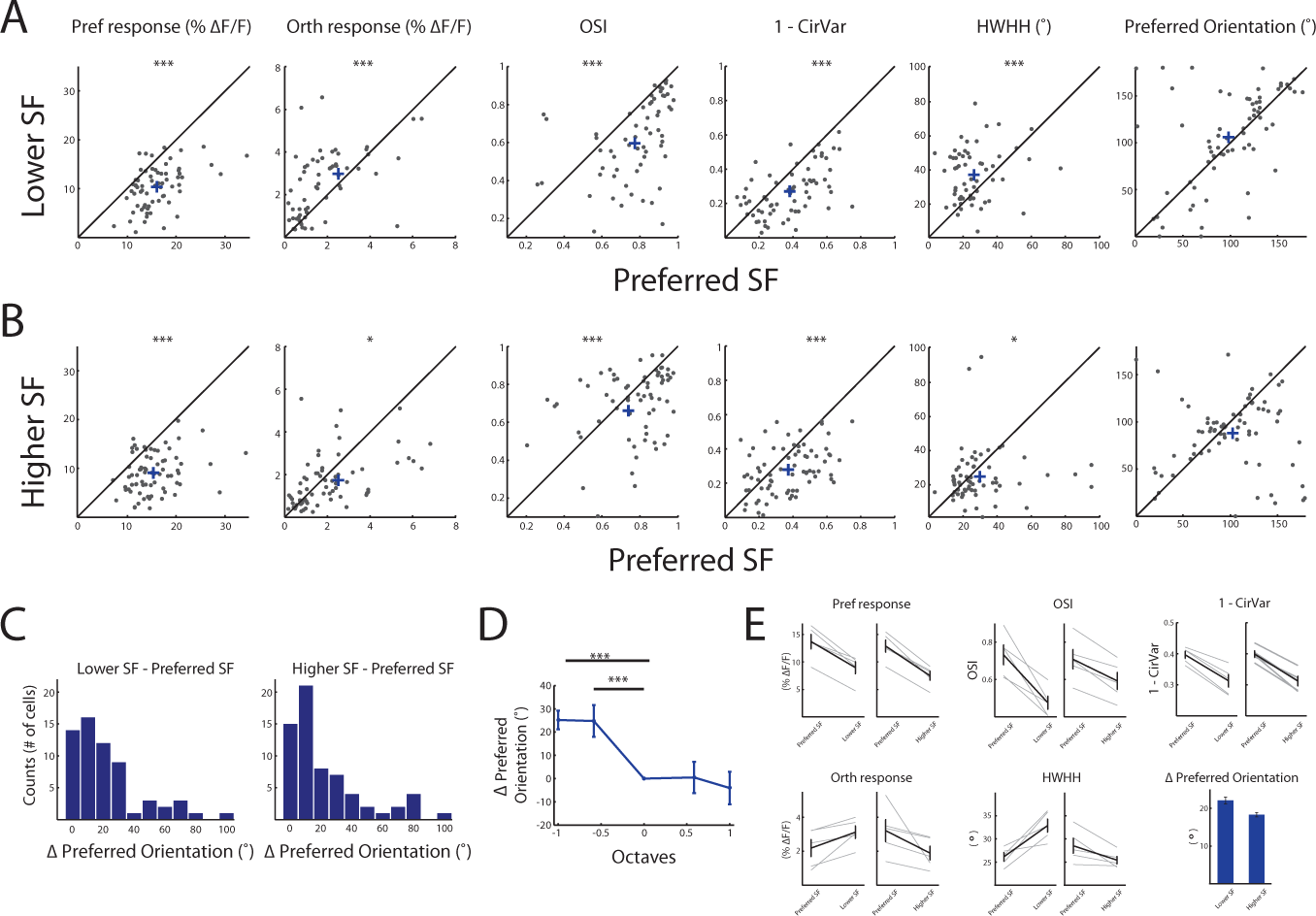
Dependence of orientation tuning curves on spatial frequency (SF) **A,** Comparison of tuning parameters between the preferred SF of each cell and one lower SF. Shown are scatter plots of cells that had preferred SF ≥ 0.02 cpd (n = 67 cells). Each circle represents a cell and cross represents population average. Shown from left to right are the response amplitude (ΔF/F) to the preferred direction (Pref), the response amplitude to the orthogonal direction (Orth), OSI, 1-CIrVar, HWHH and the preferred orientation. (Note that the preferred orientation is determined according to the average response across SFs, therefore a few cells exhibit marginally higher Pref response to non-optimal SFs). **B,** Comparison of tuning parameters between the preferred SF of each cell and one higher SF. Scatter plots of cells that had preferred SF of ≤ 0.04 cpd (n = 73 cells). Shown are the same parameters as in A. **C,** Histogram of the shift in the preferred orientation with the change in SF. Left panel shows the absolute value of the difference between the lower SF and the preferred SF and right panel shows the absolute value of the difference between the higher SF and the preferred SF. **D,** The average shift in the preferred orientation across all the tuned cells (n = 83) aligned according to the preferred SF. **E,** Population mean change in tuning curves parameters pooled from 5 animals. Gray lines depict individual animals and black line depicts the mean ± SEM across animals.

Tuning curves measured with SF *above* the preferred SF, had a decreased response to the preferred orientation (*p* < 10^−11^, n = 73 cells; Wilcoxon signed-rank test), showed a slight but significant decrease in the orthogonal-to-preferred responses (*p* < 0.01), a decrease in OSI (*p* < 0.01, Wilcoxon signed-rank test), a decrease in 1-CirVar (*p* < 10^−4^, Wilcoxon signed-rank test) and a slight decrease in HWHH (*p* < 0.05 Wilcoxon signed-rank test). Here too, we found cells with a significant shift in their preferred orientation (Fig. 2A, right panel), and indeed the Δ in preferred orientation was 18.5 ± 2.5° (mean ± SEM, Fig. 2C right panel). Further analysis of the change in the preferred orientation in non-optimal SFs revealed a monotonous averaged shift from SF *below* the preferred to SF *above* the preferred, quantified as the averaged shift across cells shown in Fig. 2C-D and exemplified in the tuning curves of single cells shown in Fig. 1D. The population results shown in Fig. 2A-B, were consistent across animals. The preferred response was decreased as expected, at lower and higher SFs (Δ4.73±0.74% and Δ5.39±0.66%, respectively; n=5 animals; 85±3.6 cells; t-test; *p*<0.005); the Orth response was increased and decreased at lower and higher SFs, respectively (Δ0.94±0.31% and Δ1.32±0.67%, respectively; t-test; *p*<0.05); the mean OSI was significantly decreased at lower and higher SFs (Δ0.26±0.05% and Δ0.12±0.02%, respectively; t-test; *p*<0.01); the mean (1-CirVAr) was significantly decreased at lower and higher SFs (Δ0.084±0.010% and Δ0.087±0.009%, respectively; t-test; *p*<0.01); the HWHH was increased and decreased at lower and higher SFs (Δ6.7±1.7% and Δ3.1±1.1%, respectively; t-test; *p*<0.05) and the preferred orientation had a mean shift of 22.1±0.9° at a lower SF and a 18.3±0.6° shift at a higher SF (mean ± SEM; Fig. 2E).

### A tilted Gabor model explains a shift in the preferred orientation at different SF

To our knowledge, the observed shift in the preferred orientation as a function of SF has not been previously reported in mice, although it has been demonstrated in cats (Jones et al., 1987). Therefore, we further investigated whether a known RF model of simple cells in V1 may explain this finding. To do so, we used a Gabor filter model with even or odd symmetry (*θ* = 0, or *θ* = π/2 respectively, where *θ* denotes the sine-wave phase, see Materials and Methods; Fig. 3A) to compute the predicted response of a neuron. A Gabor with even symmetry consists of two side-by-side antagonistic regions of equal strength whereas a Gabor with odd symmetry demonstrates a central region flanked by two antagonistic regions of equal strength (see also Fig. 4A). The neuronal response is predicted based on the dot product between the Gabor filter and the stimulus (square-wave drifting gratings). Since we imaged Ca^2+^ signals that have slow dynamics, we could not deduce the optimal phase each neuron preferred. Therefore, as the experimental tuning curves were calculated based on the maximum evoked responses, we used in our predictions the maximal response across phase (Fig. 5).

We calculated the predicted response for stimuli with various orientations and SFs, and accordingly computed the orientation tuning curves at different SFs (Fig. 3C) and examined the behavior of their parameters (Fig. 3D). First, we observed a significant difference between a Gabor with even symmetry vs. a Gabor with odd symmetry in the predicted responses to the orthogonal-to-preferred orientation. This difference also led to a difference in OSI by its definition and in 1-CIrVar. Second, and more importantly, we noticed that a symmetric Gabor, regardless of its phase, could not explain a shift in a neuron’s preferred orientation with a change in SF (Fig. 3D). Therefore, we also simulated the predicted response by altering different symmetry components of the Gabor, *e.g.* the phase of the sine wave *(θ)* and the spatial aspect ratio of the Gaussian envelope (γ; see Materials and Methods and Fig. 4A-B), but these alterations alone could not qualitatively explain a shift in the preferred orientation of the cells.

Since the classic Gabor model did not predict our experimental findings, we sought to examine alternative models. We found that one simple modification to the traditional model could explain the unexpected dependence of orientation on SF. We introduced an asymmetry by way of tilting the Gaussian envelope along the elongated axis of the 2D sine wave (Fig. 6A) and generated a filter that demonstrates a displacement of one subfield relative to the other along the RF orientation axis. We refer to this model as a tilted Gabor. Again we generated two types of tilted Gabors according to the phase of the sine wave (*θ* = 0, or *θ* = *π*/2) and then predicted the neuronal responses accordingly (Fig. 6B). We examined the dependence of orientation tuning curves on SF (Fig. 6C) and found that using this modified RF we can predict the behavior of all parameters: a change in the Orth response, a change in OSI, a change in 1-CirVar, a change in HWHH, and most importantly, a shift in the neuron’s preferred orientation. Here too we observed a difference in the preferred orientation shift between RF phases, such that RF with *θ* = 0 showed a monotonic decrease with the increase in SF whereas a tilted Gabor with *θ* = π/2 showed a non-monotonic behavior (Fig. 6D).

**Figure 6:**
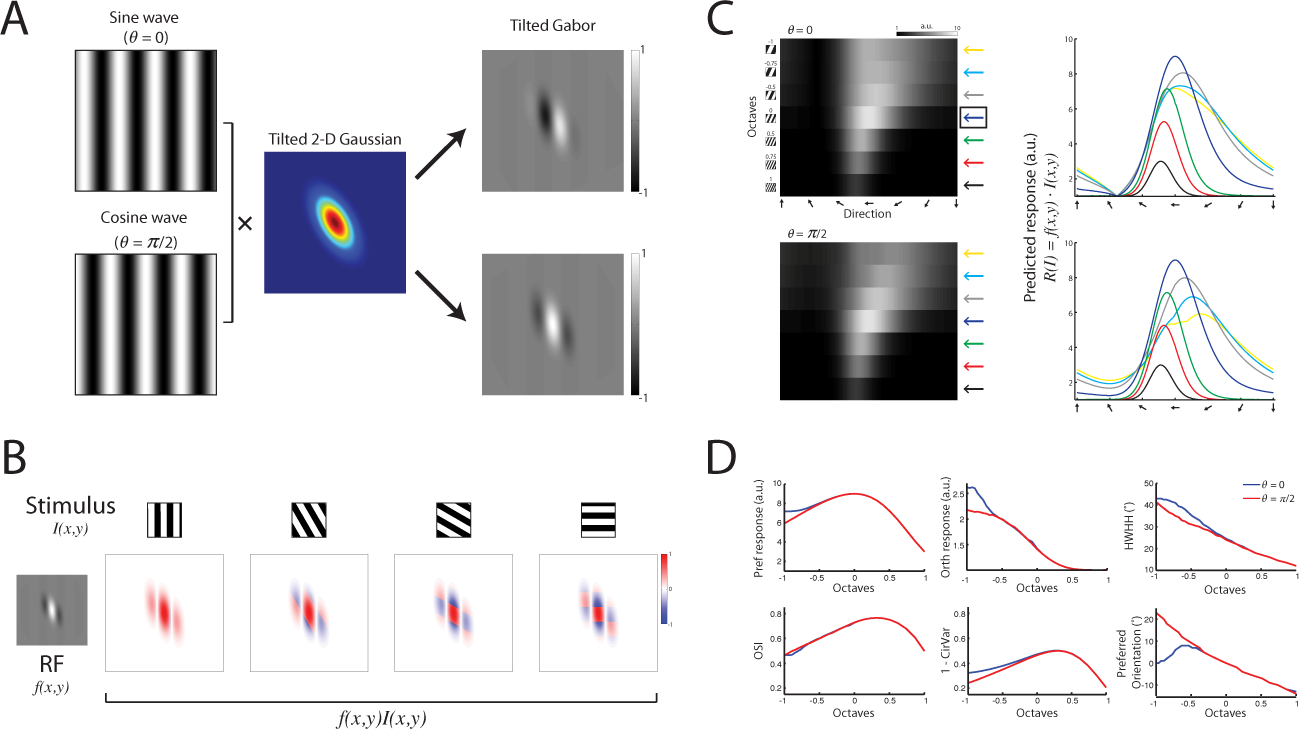
Predicting the responses of a simple cell based on RF-model of 2D tilted Gabor. **A,** A 2D tilted Gabor function: a sinusoidal plane wave weighted by a *tilted* Gaussian envelope (the Gaussian was tilted against the orientation of the sinusoidal plane wave) in two different phases: *θ* = 0 (sine wave) or *θ* = π/2 (cosine wave) shown at the top and bottom panels, respectively. **B,** Constructing spatiotemporally oriented impulse responses from gratings stimuli drifting against a *tilted* Gabor RF. Top: examples of stimuli with four various orientations. Bottom: left shows an example of a tilted Gabor RF with *θ* = π/2. On the right, 4 examples where each box depicts the overlap of the grating stimulus crossing the RF in a phase that yields the maximum response, calculated as the inner product between the stimulus and the RF. **C,** A predicted-response tuning matrix computed as the inner product between the tilted Gabor RF model shown in A, and stimuli of square-wave gratings with various orientations (1° interval) and 7 SFs. Each pixel represents the inner product between the RF and the stimulus (maximized across phase). The row marked with a blue arrow on the right denotes the preferred SF and was taken as a reference for comparing other SFs. Right: orientation tuning curves at various SFs color-coded according to the arrows shown next to the predicted tuning matrix. Top and bottom panels correspond to RF with *θ* = 0 and *0* = π/2, respectively. **D,** Comparison of orientation tuning parameters between various SFs, based on the predicted response shown in C. Blue and red lines depict the parameters calculated based on a tilted Gabor RF model with *θ* = 0 and *θ* = π/2, respectively. Shown are the response amplitude (ΔF/F) to the preferred orientation (Pref), the response amplitude to the orthogonal orientation (Orth), OSI, 1-CirVar, HWHH and the preferred orientation.

Collectively, the modified tilted Gabor model better explains our experimental data and suggests an asymmetric 2D organization of the RF On and Off subfields.

### Discussion

In this study, we measured neuronal responses from layer 2/3 of primary visual cortex of mice presented with drifting gratings of various orientations and SFs. We observed a unique dependence of orientation tuning curves on SF, which suggests that the receptive field of some cells in mouse V1 present spatial asymmetry in their RF structural organization.

### Investigating tuning curves in populations of neurons using calcium imaging

Both optical and electrical recordings techniques enable the monitoring of activity from hundreds of neurons simultaneously. Therefore, to conduct feasible experiments, one cannot tailor the repertoire of stimuli to the optimal stimulus of each and every neuron, so the number and the complexity of the stimulus are reduced. In the visual cortex, one common way to reduce the stimulus dimensionality is to measure 1D tuning curves, composed of neuronal responses evoked by presenting drifting gratings of various orientations (typically 8 or 12). This is based on the assumption that the 2D spatial structure of the receptive field can account for a large fraction of the orientation selectivity of simple cells (Lampl et al., 2001). However, orientation-tuning curves measured with a small number of points are just a 1D reduction of the neuron’s receptive field, which has a more complex structure in a high-dimensional space. And so, by this reduction we not only “pay” a *quantitative* price of reduced neural responses due to non-optimal stimulus, but we also pay a *qualitative* price, like a detection of a shift in the preferred orientation of the cells in different SFs that likely arises from an asymmetric RF structure.

### Cortical neurons have an asymmetric RF spatial structure

The RF structure of a neuron in visual cortex describes the organization of On and Off subfields in visual space, and accordingly explains which visual features a neuron is sensitive to. Simple cells in primate and cat cortex have found to present mainly 1D symmetric organization (*i.e.* even- or odd-phase symmetry along the X axis, orthogonal to the preferred orientation), though there are studies showing that cells do not demonstrate only even or odd symmetry but also exhibit to a smaller extent various phases, which breaks the 1D symmetry that exists between the relative strength of each subfield (Kulikowski and Bishop, 1981; Jones and Palmer, 1987a; Ringach, 2002).

However breaking the amplitude symmetry alone cannot explain the behavior of the observed tuning parameters (see Fig. 4). Only a 2D spatial asymmetry that introduces a shift in the relative location of the subfields can predict our experimental findings. In fact, the critical characteristic in predicting a shift in the preferred orientation is the displacement of one RF-subfield relative to the other along the orientation axis, such that the symmetry in the orthogonal axis will be broken. This 2D displacement can be easily modeled by introducing a tilt in the 2D Gaussian envelope that generates an asymmetric Gabor.

Traditionally, the Gabor filter has been proposed as a model to describe the receptive field of simple cells and successfully predicts the responses of cortical neurons of monkey and cats (Marcelja, 1980; Kulikowski et al., 1982; Daugman, 1985; Field and Tolhurst, 1986; Jones and Palmer, 1987b). The Gabor model is optimal in terms of minimizing the uncertainty associated with localizing a signal in both space and SF (Marcelja, 1980; Daugman, 1985) and therefore can create a sparse representation of natural images.

The modified model we present here keeps the basic characteristics of the classic model and introduces one additional parameter that accounts for spatial asymmetry. Although the tilted model succeeds in predicting the observed responses, there is still no experimental study in mice that mapped the cells RF directly and subsequently characterized asymmetry based on observations. However a study in cats showed that the 2D response profile of simple cells is not necessarily Cartesian Separable due to a relative displacement of the subfields, and that within a given receptive field, subfields need not be the same length (Jones and Palmer, 1987a). In addition, visual inspection of examples of cells from studies conducted in mice also revealed some displacement between the On and Off subfields of the cells’ RF (*e.g.*, Bonin et al., 2011; Ko et al., 2013; Cossell et al., 2015). However Bonin et al. used wavelets stimuli to map the cells’ RF (Selesnick et al., 2005), which are spatially symmetric by nature, while Ko et al. and Cossell et al. presented natural images but used a regularized pseudoinverse method to estimate the cells’ RF (Smyth et al., 2003), which introduces a two-dimensional smoothness constraint on the RF and might bias the RF structure towards more symmetric organization.

### Functional role of asymmetry in neuronal representations

Breaking the symmetry of the receptive field of neurons has been previously reported in the Retina and in the Hippocampus. In the Retina, asymmetry is observed between the sizes of the circular receptive fields of ON and OFF ganglion cells (Chichilnisky and Kalmar, 2002), or between the relative dendritic field size of parasol and midget cells (Dacey and Petersen, 1992), or in the asymmetric adaptation of ON and OFF ganglion cells to photopic (day) and scotopic (night) conditions (Pandarinath et al., 2010).

Hippocampal place cells exhibit an asymmetry in the spatial arrangement of their place fields. That is, as the animal moves through a cells’ receptive field, the firing rate modulates asymmetrically, where at the start of the place field the firing rate is low and at the end of the field, the firing rate is higher. The asymmetry increases as a function of familiarity (Mehta et al., 2000). This asymmetry also exists at the level of subthreshold excitatory inputs to place cells, and has been proposed to arise from a change in the balance between inhibitory inputs at the pyramidal cell soma and increases in dendritic excitation (Harvey et al., 2009). This place cell asymmetry represents a prospective coding scheme, where the increased firing skewness early in the receptive field signals upcoming place field center. The network mechanisms of this asymmetry however, are not completely known.

Furthermore, asymmetry in the visual cortex might play an important role in contour integration, in line with the Gestalt principles of perceptual grouping (Metzger, 2006), specifically the law of “good continuation” (Field et al., 1993). Since neighboring neurons share RF subfields (Smith and Hausser, 2010), then the integration of a local populations of neurons, some of which exhibit spatial RF asymmetry, may be the basis for cortical representation of spatial continuity in curved shapes.

## Acknowledgments

We thank Darcy Peterka, Julia Sable and Yeonsook Shin for technical support. This study was supported by Marie Curie IOF (to I.A.), Canadian Institute for Health Research (to J.J.), the NEI (DP1EY024503, R01EY011787) and DARPA SIMPLEX N66001-15-C-4032. This material is based upon work supported by, or in part by, the U.S. Army Research Laboratory and the U.S. Army Research Office under contract number W911NF-12-1-0594 (MURI).

## References

Andrews BW, Pollen DA (1979) Relationship between spatial frequency selectivity and receptive field profile of simple cells. J Physiol 287:163–176.

Bonin V, Histed MH, Yurgenson S, Reid RC (2011) Local diversity and fine-scale organization of receptive fields in mouse visual cortex. J Neurosci 31:18506–18521.

Brainard DH (1997) The Psychophysics Toolbox. Spat Vis 10:433–436.

Chichilnisky EJ, Kalmar RS (2002) Functional asymmetries in ON and OFF ganglion cells of primate retina. J Neurosci 22:2737–2747.

Cossell L, Iacaruso MF, Muir DR, Houlton R, Sader EN, Ko H, Hofer SB, Mrsic-Flogel TD (2015) Functional organization of excitatory synaptic strength in primary visual cortex. Nature 518:399–403.

Dacey DM, Petersen MR (1992) Dendritic field size and morphology of midget and parasol ganglion cells of the human retina. Proc Natl Acad Sci U S A 89:9666–9670.

Daugman JG (1985) Uncertainty relation for resolution in space, spatial frequency, and orientation optimized by two-dimensional visual cortical filters. Journal of the Optical Society of America A, Optics and image science 2:1160–1169.

Field DJ, Tolhurst DJ (1986) The structure and symmetry of simple-cell receptive-field profiles in the cat’s visual cortex. Proceedings of the Royal Society of London Series B, Biological sciences 228:379–400.

Field DJ, Hayes A, Hess RF (1993) Contour integration by the human visual system: evidence for a local “association field”. Vision research 33:173–193.

Gabor D (1946) Theory Of Communication. J IEE (London) 93:429–457.

Harvey CD, Collman F, Dombeck DA, Tank DW (2009) Intracellular dynamics of hippocampal place cells during virtual navigation. Nature 461:941–946.

Hubel DH, Wiesel TN (1962) Receptive fields, binocular interaction and functional architecture in the cat’s visual cortex. J Physiol 160:106–154.

Jones JP, Palmer LA (1987a) The two-dimensional spatial structure of simple receptive fields in cat striate cortex. J Neurophysiol 58:1187–1211.

Jones JP, Palmer LA (1987b) An evaluation of the two-dimensional Gabor filter model of simple receptive fields in cat striate cortex. J Neurophysiol 58:1233–1258.

Jones JP, Stepnoski A, Palmer LA (1987) The two-dimensional spectral structure of simple receptive fields in cat striate cortex. J Neurophysiol 58:1212–1232.

Kerlin AM, Andermann ML, Berezovskii VK, Reid RC (2010) Broadly tuned response properties of diverse inhibitory neuron subtypes in mouse visual cortex. Neuron 67:858–871.

Kleiner M, Brainard D, Pelli D (2007) “What’s new in Psychtoolbox-3?“ Perception 36:ECVP Abstract Supplement.

Ko H, Cossell L, Baragli C, Antolik J, Clopath C, Hofer SB, Mrsic-Flogel TD (2013) The emergence of functional microcircuits in visual cortex. Nature 496:96–100.

Kulikowski JJ, Bishop PO (1981) Linear analysis of the responses of simple cells in the cat visual cortex. Exp Brain Res 44:386–400.

Kulikowski JJ, Marcelja S, Bishop PO (1982) Theory of spatial position and spatial frequency relations in the receptive fields of simple cells in the visual cortex. Biological cybernetics 43:187–198.

Lampl I, Anderson JS, Gillespie DC, Ferster D (2001) Prediction of orientation selectivity from receptive field architecture in simple cells of cat visual cortex. Neuron 30:263–274.

Marcelja S (1980) Mathematical description of the responses of simple cortical cells. Journal of the Optical Society of America 70:1297–1300.

Marshel JH, Garrett ME, Nauhaus I, Callaway EM (2011) Functional specialization of seven mouse visual cortical areas. Neuron 72:1040–1054.

Mazurek M, Kager M, Van Hooser SD (2014) Robust quantification of orientation selectivity and direction selectivity. Front Neural Circuits 8:92.

Mehta MR, Quirk MC, Wilson MA (2000) Experience-dependent asymmetric shape of hippocampal receptive fields. Neuron 25:707–715.

Metzger W (2006) Laws of seeing. Cambridge, Mass.: MIT Press.

Miller JE, Ayzenshtat I, Carrillo-Reid L, Yuste R (2014) Visual stimuli recruit intrinsically generated cortical ensembles. Proc Natl Acad Sci U S A 111:E4053–4061.

Niell CM, Stryker MP (2008) Highly selective receptive fields in mouse visual cortex. J Neurosci 28:7520–7536.

Nimmerjahn A, Kirchhoff F, Kerr JN, Helmchen F (2004) Sulforhodamine 101 as a specific marker of astroglia in the neocortex in vivo. Nature methods 1:31–37.

Pandarinath C, Victor JD, Nirenberg S (2010) Symmetry breakdown in the ON and OFF pathways of the retina at night: functional implications. J Neurosci 30:10006–10014.

Pelli DG (1997) The VideoToolbox software for visual psychophysics: transforming numbers into movies. Spat Vis 10:437–442.

Piscopo DM, El-Danaf RN, Huberman AD, Niell CM (2013) Diverse visual features encoded in mouse lateral geniculate nucleus. J Neurosci 33:4642–4656.

Ringach DL (2002) Spatial structure and symmetry of simple-cell receptive fields in macaque primary visual cortex. J Neurophysiol 88:455–463.

Ringach DL, Shapley RM, Hawken MJ (2002) Orientation selectivity in macaque V1: diversity and laminar dependence. J Neurosci 22:5639–5651.

Selesnick IW, Baraniuk RG, Kingsbury NC (2005) The dual-tree complex wavelet transform. IEEE Signal Processing Magazine 22:123–151.

Skottun BC, De Valois RL, Grosof DH, Movshon JA, Albrecht DG, Bonds AB (1991) Classifying simple and complex cells on the basis of response modulation. Vision research 31:1079–1086.

Smith SL, Hausser M (2010) Parallel processing of visual space by neighboring neurons in mouse visual cortex. Nat Neurosci 13:1144–1149.

Smyth D, Willmore B, Baker GE, Thompson ID, Tolhurst DJ (2003) The receptive-field organization of simple cells in primary visual cortex of ferrets under natural scene stimulation. J Neurosci 23:4746–4759.

Thevenaz P, Ruttimann UE, Unser M (1998) A pyramid approach to subpixel registration based on intensity. IEEE Trans Image Process 7:27–41.

Vidyasagar TR, Siguenza JA (1985) Relationship between orientation tuning and spatial frequency in neurones of cat area 17. Exp Brain Res 57:628–631.

Webster MA, De Valois RL (1985) Relationship between spatial-frequency and orientation tuning of striate-cortex cells. Journal of the Optical Society of America A, Optics and image science 2:1124–1132.

Zhu W, Xing D, Shelley M, Shapley R (2010) Correlation between spatial frequency and orientation selectivity in V1 cortex: implications of a network model. Vision research 50:2261–2273.

